# Recurrent neural networks learn robust representations by dynamically balancing compression and expansion

**DOI:** 10.1101/564476

**Authors:** Matthew Farrell, Stefano Recanatesi, Timothy Moore, Guillaume Lajoie, Eric Shea-Brown

## Abstract

Recordings of neural circuits in the brain reveal extraordinary dynamical richness and high variability. At the same time, dimensionality reduction techniques generally uncover low-dimensional structures underlying these dynamics. What determines the dimensionality of activity in neural circuits? What is the functional role of this dimensionality in behavior and task learning? In this work we address these questions using recurrent neural network (RNN) models, which have recently shown promise in predicting and explaining brain dynamics. Through simulations and mathematical analysis, we show how the dimensionality of RNN activity evolves over time and over stages of learning. We find that RNNs can learn to balance tendencies to expand and compress dimensionality in a way that matches task demands and further generalizes to new data. Strongly chaotic networks appear particularly adept in learning this balance in the case of classifying low-dimensional inputs, combining the natural tendency of chaos to expand dimensionality with opportunistic compression driven by stochastic gradient descent to form representations with good generalization properties. These findings shed light on fundamental dynamical mechanisms by which neural networks solve tasks with robust representations that generalize to new cases.

## INTRODUCTION

Dynamics shape computation in brain circuits. These dynamics arise from highly recurrent and complex networks of interconnected neurons, and neural trajectories observed in cortical areas are correspondingly rich and variable across stimuli, time, and trials. Despite this high degree of variability in neural responses, repeatable and reliable activity structure is often unveiled by dimensionality reduction procedures [15, 47]. Rather than being set by, say, the number of neurons in the circuit, the *effective* dimensionality of the neural activity (or neural “representation”) seems to be intimately linked to the complexity of the function, or behavior, that the neural circuit fulfills or produces [14, 19, 50, 57]. Similar task-dependent trends in dimensionality can manifest in artificial networks used in machine learning systems trained using optimization algorithms (e.g., [14, 42, 62]). Bridging between machine learning and neuroscience, artificial networks are powerful tools for investigating dynamical representations in controlled settings, and enable tests of theoretical hypotheses that can be leveraged to formulate experimental predictions (reviewed in [6]).

The connection between task complexity and representation dimension is especially intriguing in light of fundamental ideas in learning theory. On the one hand, high-dimensional representations can subserve complex and general computations that nonlinearly combine many features of inputs [8, 13, 18, 39, 52, 61]. On the other, low-dimensional representations that preserve only essential features needed for specific tasks can allow learning based on fewer parameters and examples, and hence with better “generalization” (for reviews, see [7, 9, 18, 60], and see [10, 11] for important steps toward characterizing the coding properties of general representation geometries).

What are the mechanisms that regulate the effective dimensionality of network activity across many trials and inputs? While dimensionality modulation can occur due to hard structural constraints such as “bottleneck” layers with a small number of neurons [23], here we consider the case where the number of neurons is not a constraining factor. We consider two factors: chaos and learning. Frequently encountered in recurrent neural network (RNN) models of brain function, dynamical chaos (whereby tiny changes in internal states are amplified by unstable, but deterministic, dynamics) provides a parsimonious explanation for both repeatable structure as well as internally generated variability seen in highly recurrent brain networks such as cortical circuits [17, 30, 38, 40, 55, 56, 59, 63]. In the reservoir computing framework, chaos in an RNN can increase the diversity of patterns and dimension of activity the network produces through time in response to inputs, in some cases increasing the system’s computational abilities [26, 35, 41]. While chaos-driven expansion as determined by fixed recurrent weights has been explored in simulations [35], the attributes of this phenomenon as recurrent weights evolve through training are less understood (but see [16, 58]).

The goal of the current work is to elucidate how an RNN’s representation and dynamics are shaped by learning to balance the two desiderata of compressing and expanding the dimension of input data. Notably, we find evidence that stochastic gradient descent (SGD) is predisposed to compress dimension, even when expansion is necessary to solve the task. Chaos in our case provides a key ingredient for effective learning: directions of expansion that coexist with many directions of compression. SGD is often effective at making use of this expansion while still compressing opportunistically without sacrificing task performance. This results in learned representations that balance expansion and compression and that consequently lend themselves to good generalization. The same balance of expansion and compression in chaotic networks may account for the highly variable nature of neural activity, and suggests that further optimizing this trade-off may be a method for improving the flexibility and accuracy of artificial neural networks. Finally, we analyze how SGD compresses dimensionality in neural networks. Our work is similar to the information compression phenomenon explored in [53, 54], but instead focusing explicitly on the geometry of the representation. In addition, while the noise generated by SGD in [53] is treated as additive and Gaussian, here we consider the true noise generated by two steps of SGD in a more simplified model. This contributes to revealing why this ubiquitous algorithm often finds solutions that generalize (e.g. [2, 4, 20, 29, 33, 36, 53, 54]).

## RESULTS

### Model and task overview

We investigate the dynamics of recurrent networks learning to classify static inputs. Our network model is based on standard RNN models used in machine learning [22, 37]. Interactions between *N* = 200 neural units are determined by a randomly initialized recurrent connectivity matrix, and unit activations pass through a hyperbolic tangent nonlinearity. The dynamics of the network initialized in this way can be modulated from stable to chaotic by increasing the average magnitude of the initial neural coupling strength, and vice-versa. We compare chaotic networks initialized near the transition point to chaos (said to be on the “edge of chaos”), to “strongly chaotic” networks well past the transition point, as they learn to solve the task described below. To measure chaos, we numerically compute the *Lyapunov exponents* of the system (see Supplemental Information for more details).

Inputs are *N*-dimensional constant vectors. They are selected from Gaussian-distributed clusters whose means are distributed randomly within a *d*-dimensional, random subspace of the *N*-dimensional neural activity space. We call this subspace the input’s *ambient space* and consider two scenarios for the ambient dimension: *d* = 2 and *d* = *N*. Each cluster is assigned one of two class labels, at random, as illustrated in Fig. 1. While this schematic only shows six clusters for clarity, in our simulations we use 60 clusters. Each cluster is assigned one of two class labels: 30 clusters for label 1, and 30 for label 2. The task proceeds as follows: (1) a random input is selected from one of the clusters and presented to the network for one timestep; (2) the network’s dynamics evolve undriven during a delay period of 9 timesteps; (3) a linear readout of the network’s state at timestep 10 is used to classify the input into one of the two classes. Recurrent and output weights are adjusted via an SGD routine to minimize a cross entropy loss function, and classification performance is evaluated on novel inputs not used for training. We find our results qualitatively robust to moderate changes in the choice of model parameters, as well as using squared error loss instead of cross entropy. See Fig. S4, where we train our network on the MNIST digit classification database [34].

**FIG. 1.**
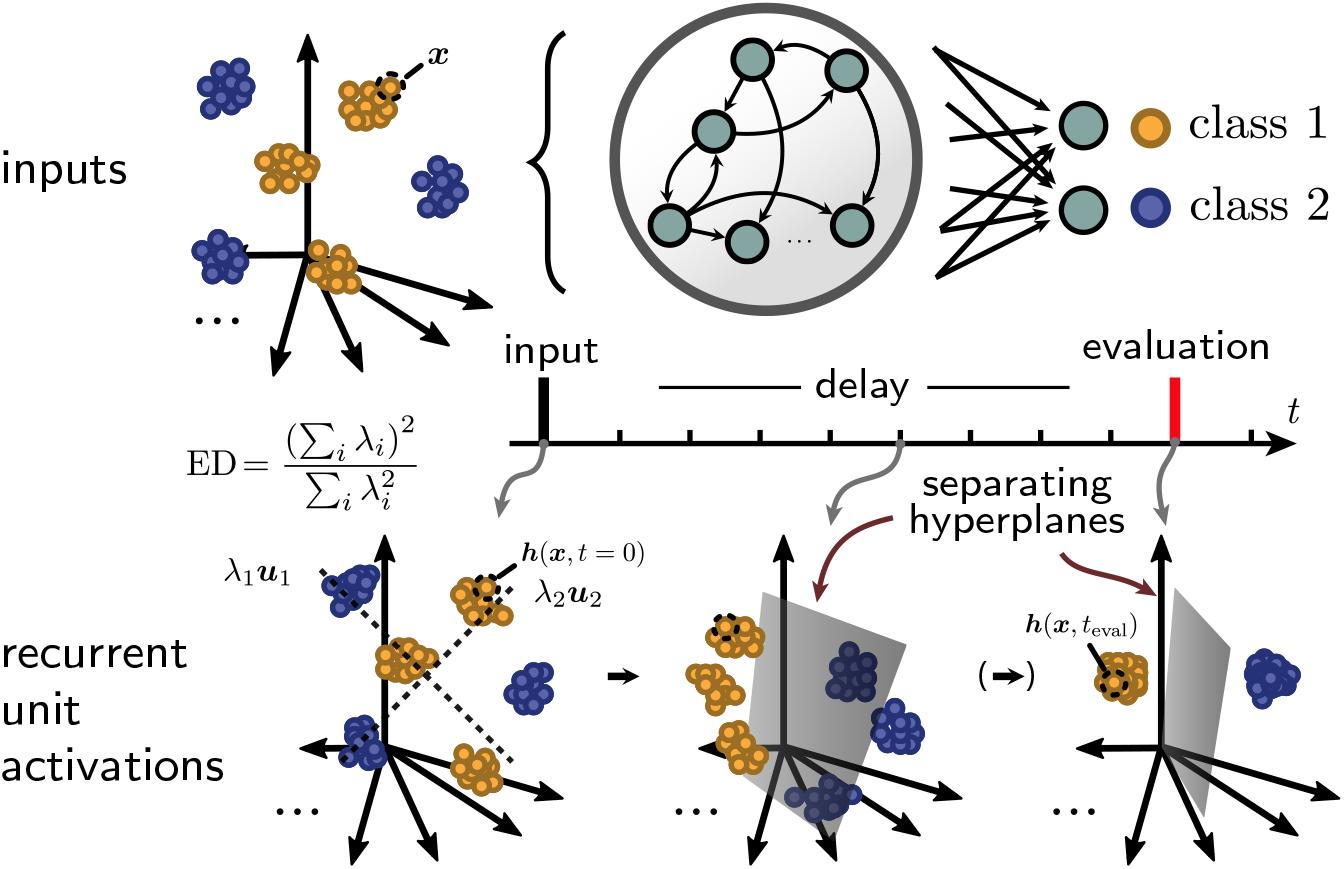
Task and model schematic. Input clusters are distributed in a *d*-dimensional subspace of neural space. An input ***x*** is shown to the network for one timestep, and after a delay period in which the network evolves undriven, the network state ***h***(***x***, *t*_eval_) is linearly read out at a timepoint *t*_eval_ in order to classify the input. Bottom: Schematic of the response of the network to the ensemble of inputs, viewed at snapshots in time. Through training, the network attempts to form a representation that allows a separating hyperplane. The network may also bring points belonging to the same class together to form a more compact representation. Bottom left: Dashed lines depict the top two eigenvectors of the covariance of the representation, scaled by the corresponding eigenvalues. To capture the degree to which activations fill space, we use the effective dimensionality (ED), which measures the participation ratio of all of the eigenvalues.

### SGD trains networks to compress high-dimensional inputs

We start by considering the classification of high-dimensional inputs, where the input ambient space dimensionality *d* is equal to the number *N* of recurrent units (see Fig. 2a for a visualization). While classification in networks is often viewed from the perspective of making inputs linearly separable, in our scenario the data are already linearly separable in the input space. This focuses our attention on what, if anything, networks learn to do beyond producing linear separability.

**FIG. 2.**
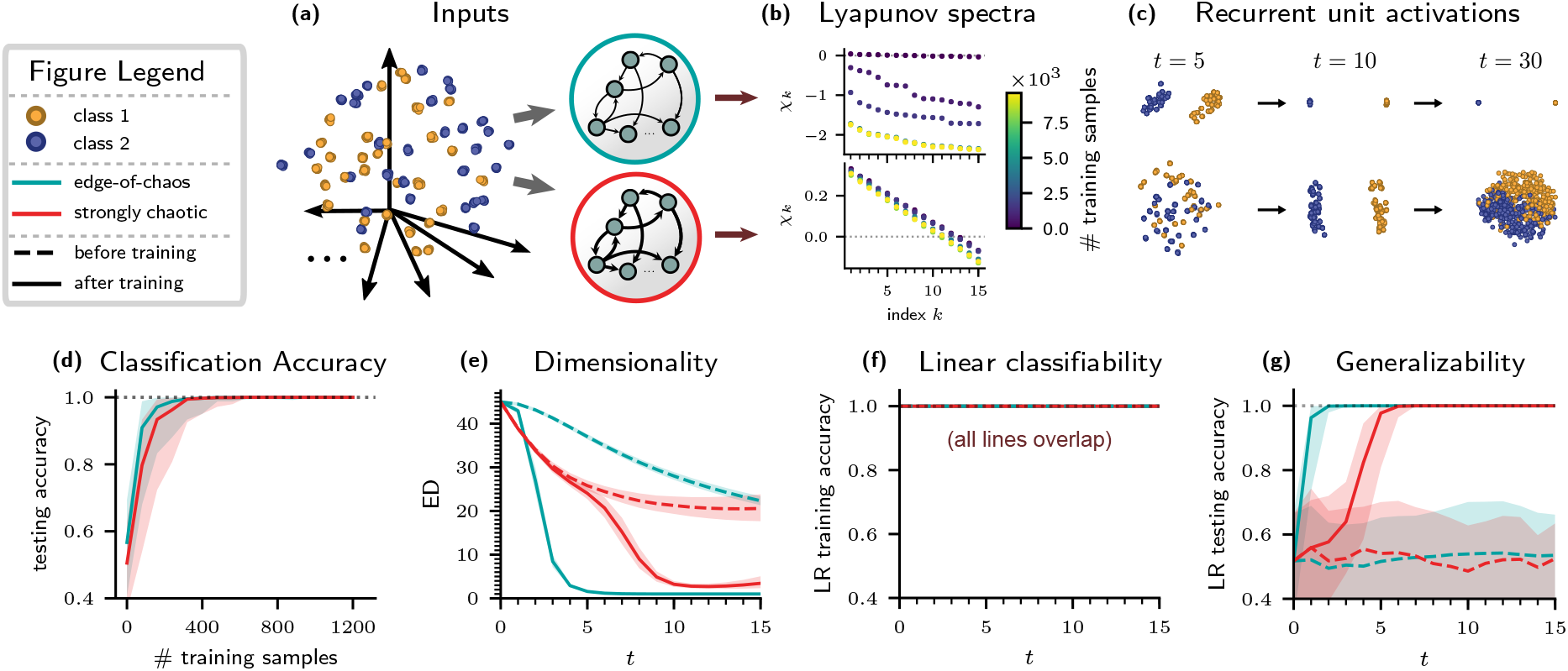
Comparison of edge-of-chaos (blue-green) and strongly chaotic (red) networks classifying high-dimensional inputs. Input color (yellow-orange or purple) denotes true class label. Shaded regions indicate 75% probability mass of gamma distributions fit to values of the dependent variables over a sample of 30 network and input realizations, with solid lines indicating medians. “After training” designates networks that have been trained on 9,600 input samples. (a) Schematic of the task. (b) Lyapunov exponents measured through training. Each plot is of a single network and input realization. Error bars (too small to see) denote standard error of the mean. Top: Edge-of-chaos network. Bottom: strongly chaotic network. (c) Activations of recurrent units responding to an ensemble of 600 inputs, plotted as “snapshots” in time in principal component space. Top: Edge-of-chaos network. Bottom: strongly chaotic network. (d) Testing accuracy measured through training. (e) Effective dimensionality (ED) of the network representation through time *t*. (f) Mean accuracy of a logistic regression (LR) linear classifier trained on the recurrent unit activations at each timepoint *t*. (g) Mean testing accuracy of an LR classifier. The model was trained on data drawn from 80% of the input clusters (816 input points) and tested on held-out data from the remaining 20% of input clusters (204 input points).

We study networks in two dynamical regimes: a strongly chaotic network and one that is initialized at the edge of chaos. Fig. 2b measures the degree of chaos of the edge-of-chaos and strongly chaotic regimes by plotting the top 15 Lyapunov exponents *χ_k_* of the networks through training. In these plots, positive values of *χ_k_* indicate chaotic dynamics. If all *χ_k_* are negative, then the dynamics are non-chaotic (stable), meaning that trajectories converge to stable fixed points or stable limit cycles. The edge-of-chaos network is weakly chaotic before training and becomes stable after training, while the strongly chaotic network is chaotic both before and after training. While each of the plots in Fig. 2b only shows the exponents for a single network realization and a particular positioning of input clusters, they capture the general qualitative behavior of the two dynamical regimes for *d* = *N*.

As shown in Fig. 2d, both networks easily achieve perfect testing accuracy on the delayed classification task. What is then of interest is how properties of the network’s *internal representation* change over the course of training. In Fig. 2c we plot the top two principal components of the network responses to 600 input samples at snapshots in time. The top row corresponds to the edge-of-chaos network, where we see that points belonging to different classes are pulled apart while points within the same class are compressed together in all dimensions (see Fig S1 for a quantification of this phenomenon). The case is similar for the strongly chaotic network (bottom row), except that the classes begin to mix back together in the top two PCs after the evaluation time *t*_eval_ = 10. Our analysis of the Lyapunov exponents indicates that the edge-of-chaos network forms fixed points that persistently solve the task, while the strongly chaotic network forms chaotic attractors that transiently solve the task.

This compression phenomenon is partly captured by measuring the *effective dimensionality* (ED) of the representation [27, 48]. The equation for the ED of a set of *S* points ***V*** = [*v_si_*] ∈ ℝ^*S* × *d*^ with ambient dimension *d* is

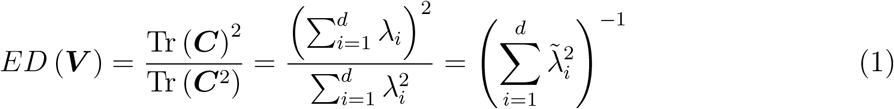

where *C_ij_* = 〈*v_si_v_sj_*〉_*s*_ – 〈*v_si_*〉_*s*_ 〈*v_sj_*〉_*s*_ is the covariance matrix of ***V*** and the 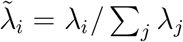 are the normalized eigenvalues of ***C***. A visual intuition for this quantity is shown in Fig. 1. ED can be roughly thought of as the number of principal components needed to capture most of the variance of the data [19].

In Fig. 2e the ED is plotted through time *t*. For each time point *t* the matrix ***V*** = ***V***(*t*) used to compute ED has dimensions *S* × *N* where *S* is the number of input samples shown to the network. The EDs of the trained networks are approximately equal to that of the input at time *t* = 0, since the initial states only differ from the inputs by one application of the nonlinearity (see Methods). The dimensionality drops with increasing *t* and is highly compressed at the evaluation time *t*_eval_ = 10. This compression results both from increasing distances between different classes as well as decreasing distances within classes (see Fig. 2c and Fig S1). The degree of these trends depends somewhat on the learning procedure, with more aggressive weight updates resulting in faster dimensionality compression (data not shown). However, the general behavior is robust to moderate changes in the learning algorithm parameters. This compression can be viewed through the lens of building *invariance* in the network representation [1, 20, 21], in this case invariance to input cluster identity (data not shown).

We next study the coding properties of these representations. Fig. 2f shows the mean accuracy of a logistic regression (LR) linear classifier trained to classify the network representation at each timepoint. This confirms the point made above: the input data is linearly separable, and this property is retained by the network dynamics. The interesting properties of the network computation lie elsewhere, in that the learned dynamics and resulting dimensionality compression lead to better generalization properties of the representation. In Fig. 2g, we measure generalization by first training an LR classifier on the network response to inputs drawn from a fixed 80% of the input clusters, and then measuring the accuracy of this classifier on the network response to samples drawn from the remaining 20% of the clusters. The dashed lines indicate that, while the representations in the untrained networks are linearly separable, a linear classifier trained on these representations does not generalize well to held-out clusters. In contrast, after training, the network representations become increasingly generalizable through time *t*, eventually allowing for perfect classification accuracy on held out clusters and maintaining this for future times.

We emphasize that in this case linear separability – the property needed to solve the task with perfect accuracy – requires neither dimensionality expansion nor compression of inputs. Nevertheless, we find that networks **do** learn to strongly compress their inputs, at the same time achieving representations that lend themselves to good generalization.

### Chaos drives dimensionality expansion, enabling SGD to find a balance between expansion and compression

We next turn our attention to inputs embedded in a two-dimensional ambient space, *d* = 2 (see Fig. 3a for a visualization). In this case, the two classes are generally far from being linearly separable (classification boundaries must be curved and nonlinear to separate the 60 clusters randomly distributed in two-dimensional space). As a consequence, it is difficult for the network to classify without first increasing the dimensionality of its representation.

**FIG. 3.**
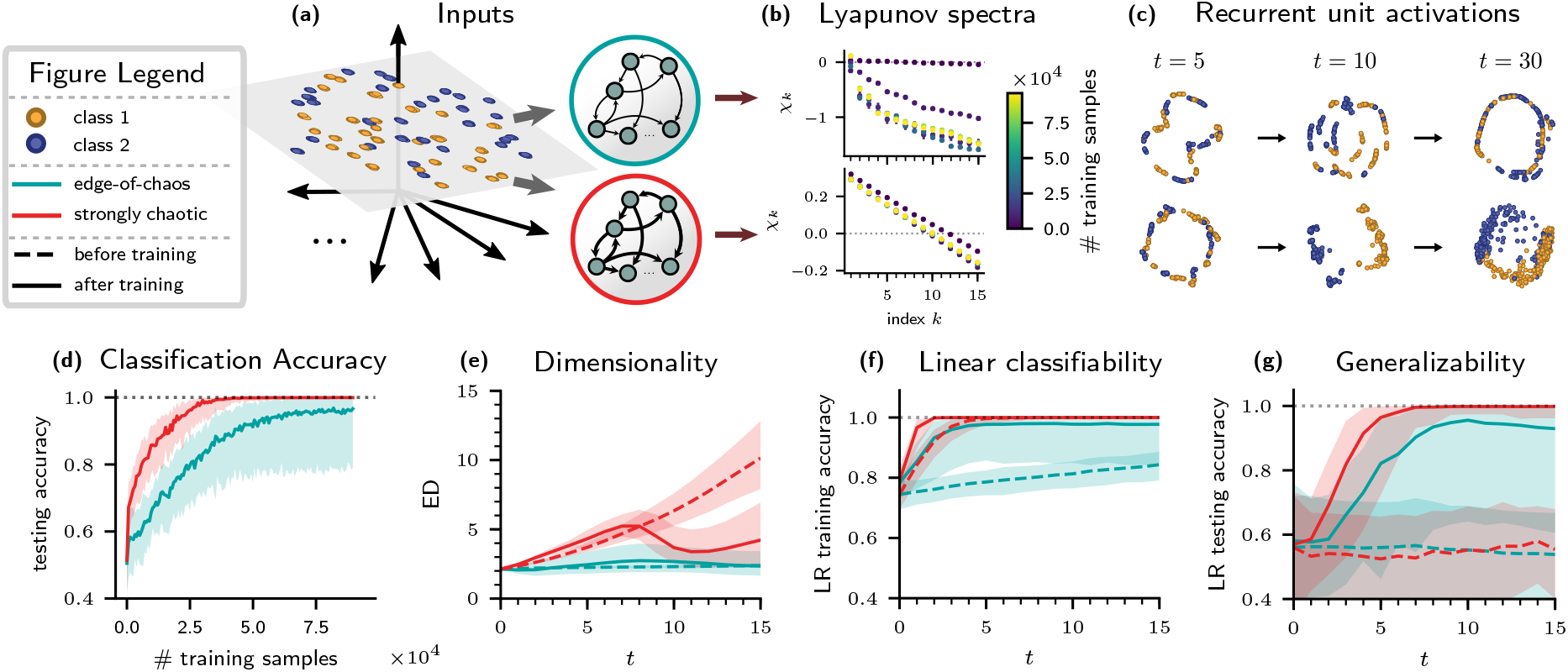
Comparison of edge-of-chaos and strongly chaotic networks classifying low-dimensional inputs. Details are similar to Fig. 2. “After training” designates networks that have been trained on 96,000 input samples. (a) Inputs lie on a two-dimensional plane cutting through recurrent unit space. (b-g) Descriptions as in Fig. 2.

Fig. 3 compares the behavior of the edge-of-chaos and strongly chaotic networks trained on this task. In this case, the edge-of-chaos network remains on the edge-of-chaos through training (Fig. 3b). The strongly chaotic network remains strongly chaotic during training. In comparing the representations in Fig. 3c, we find indications that the strongly chaotic network (bottom) is more successful at achieving class separation at time *t*_eval_ = 10 than the edge-of-chaos network (top). Fig. 3d confirms that this is indeed the case: the strongly chaotic network learns to achieves near-perfect classification accuracy, while the edge-of-chaos network is not as successful (Fig. 3d).

In looking at the dimensionality of the trained networks through time (Fig. 3e), we find that both initially expand dimensionality until about *t* = 7, with the expansion being much more dramatic for the strongly chaotic network (solid lines). The strongly chaotic network then reverses course to compress dimensionality up to time *t*_eval_ = 10. The dimensionality expansion of the strongly chaotic network up to *t* = 7 seems to follow from its natural tendency to expand dimensionality before training, in contrast to the edge-of-chaos network (dashed lines). The compression from *t* = 7 to *t* = 10 is then learned through training.

In Fig. 3f we see that the strongly chaotic network creates a linearly separable representation both before and after training, while the representations for the edge-of-chaos network are far from linearly separable before training. This is consistent with the expansion of dimensionality that occurs even before training in the strongly chaotic network. The generalization of both networks improves with training (Fig. 3g), with the strongly chaotic network achieving better generalization performance.

We next turn our attention to a more challenging task, where input is only shown to a subset of neurons (Fig. 4a); here, two. In this case, dimensionality expansion can have the added benefit of enlisting more neurons to aid in the computation (e.g. [31]). We find that the strongly chaotic network is still able to solve the task near-perfectly, while the edge-of-chaos network’s median performance remains at about 75% (Fig. 4c). Looking at ED, we find that the edge-of-chaos network does not learn to significantly expand ED, while the strongly chaotic network is higher-dimensional before training and learns to expand its dimensionality even further (Fig. 4e). This expansion again results in good linear separability by the evaluation time *t* = 10, even before training (Fig. 4f).

**FIG. 4.**
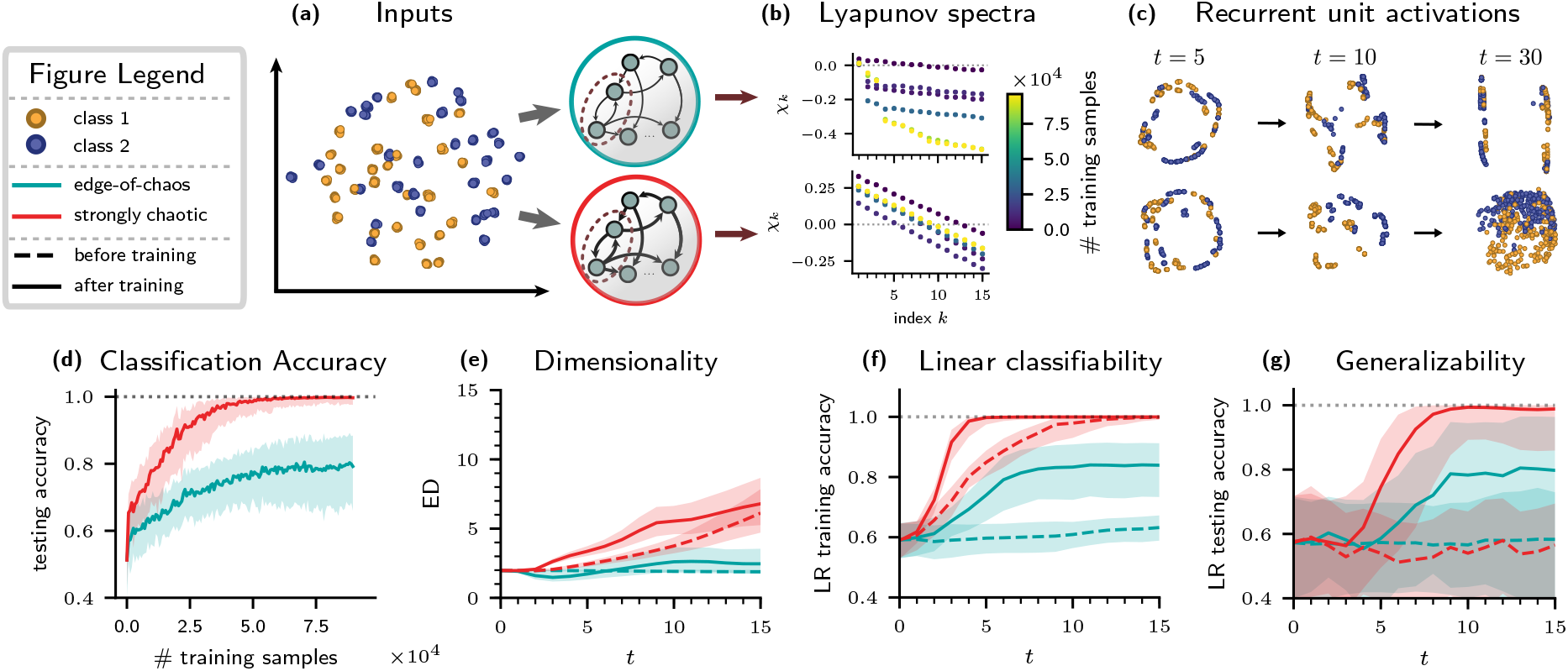
Comparison of edge-of-chaos and strongly chaotic networks classifying two-dimensional inputs fed to two neurons. Details are similar to Figs. 2 and 3. “After training” designates networks that have been trained on 96,000 input samples. (a) Schematic of the task. Two-dimensional input is delivered to two randomly selected neurons in the network. (b-g) Descriptions as in Fig. 2.

The generalization properties of the strongly chaotic network’s representation improve after training, even though dimensionality compression does not occur (Fig. 4g, solid red line). However, comparing this network at the evaluation time with the strongly chaotic network trained on distributed inputs (Fig. 3g, solid red line), we find that here the generalization score is less reliable, with larger error bars. This suggests that the dimensionality compression exhibited in Fig. 3e, solid red line, plays a role in improving generalization properties, while constrained dimensionality expansion as in Fig. 4e, solid red line, can still allow for relatively good (albeit not as good) generalization. Indeed, while the strongly chaotic network expands dimensionality without a clear compression phase, the dimensionality is still small at the evaluation time relative to the size of the network.

### Mechanistic explanations for compression

Our networks exhibit dimensionality compression when tasked with classifying highdimensional, linearly separable inputs. Above we have discussed the benefits of this compression from the perspective of generalization; in this section, we give mechanistic explanations for why this compression occurs. More specifically, for gradient descent parameter updates, we show how noise can can lead to weight changes that compress the dimensionality of internal unit representations, and then apply this analysis to understand how SGD drives compression. See [25] for a more thorough analysis in the case of networks trained by a particular unsupervised learning rule, [28] for compression induced by the “Pseudo-Inverse” learning rule, and [49] for a recently developed alternative approach to analyzing compression induced by SGD. Our result is related to the pioneering idea of information compression which is studied in, e.g.([53, 54]), which implicates the same driving mechanism (gradient noise) and establishes the resulting impact on generalization.

To shed light on the compression behavior illustrated in Figs. 2d, 2e, 3c and 3e – where points that belong to the same class are brought close together by the dynamics of the trained network – we show how compression can occur in a simplified scenario. In particular, we consider a linear, single-hidden-layer feedforward network. Our analysis proceeds in two steps. First, we show how gradient updates with isotropic noise injected in the output weights leads to compression in all directions orthogonal to the readouts. This suggests a crucial role for variability in driving compression. Second, we show how the noisy variability of the network weights generated by SGD can also result in compression orthogonal to the readout direction.

In a single-layer linear network, the internal neural activations responding to a single input sample ***x*** are ***h***(***x***) = ***Wx*** + ***b***, and the scalar output is 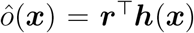. Since significant compression occurs for both cross entropy as well as squared error loss in our simulations, for ease of analysis we consider squared error loss 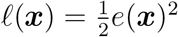, where 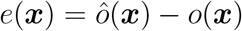 and *o* maps the input ***x*** to its corresponding class label, either +1 or −1. The gradient descent learning updates ***W*** ← ***W*** + *δ****W*** and ***b*** ← ***b*** + *δ****b*** over a batch of input samples *B* are:

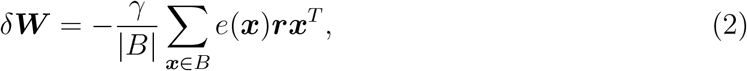

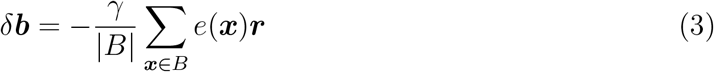

Here *γ* is the *learning rate* and ***rx***^*T*^ is the outer product of ***r*** and ***x***. The update rule results in a corresponding update ***h*** ← ***h*** + *δ****h*** where *δ****h***(***x***′) = *δ****Wx***′ + *δ****b***.

Note that the change of hidden representation *δ****h*** points in the direction of the readout vector: *δ****h***(***x***′) ∝ ***r***. If output weights ***r*** are unchanged by the update rule, then gradient descent parameter updates can modulate the response only along this single dimension. This modulation can be expected to cause compression in this single dimension when gradient descent reduces the loss, as the average loss is minimized when ***r***^⊤^***h***(***x***′) = *o*(***x***′) for all ***x***′. Satisfying this equation implies that ***h***(***x***′) for *o*(***x***′) = +1 has minimized its variation along ***r***, and similarly for *o*(***x***′) = −1.

To see reduction of variability in all dimensions like that exhibited by Fig. 2f, rather than only along a single dimension, we need to consider how the direction of compression can change across learning. We turn to this next. For all subsequent calculations, we assume that inputs have zero mean and covariance 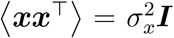 (note that this is approximately satisfied by the inputs used for the simulations above).

Let ***h***_*k*_ denote the hidden representation after *k* steps of gradient descent, and similarly for parameters ***W, b***, and ***r***. A single step of gradient descent results in the representation ***h***_1_(***x***′) = ***h***_0_(***x***′) + *δ****W***_0_***x***′ + *δ****b***_0_. Averaging over batches *B*, this becomes

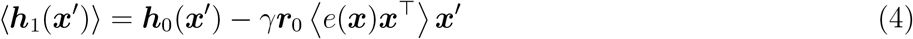

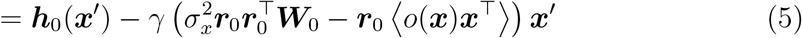

We will first consider variability in the initial readout direction ***r***_0_ by modeling it as isotropic additive noise: 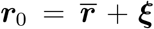 where ***ξ*** is a random variable with zero mean and covariance 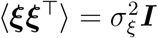 and 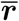 is fixed. This noise is a simple model of variability that could arise, for instance, from previous steps of SGD. Let 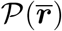 denote any vector proportional to 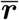 (this notation functions similarly to big O notation). These terms contribute only along a single dimension, and we can expect compression in this direction as long as the loss is decreasing (see discussion above), so we don’t analyze these terms further here and instead focus on the components orthogonal to 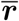. We then obtain

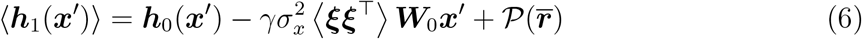

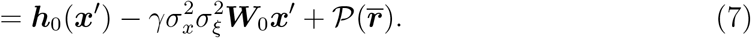

If we further assume that the bias is aligned with the readout, 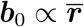, then

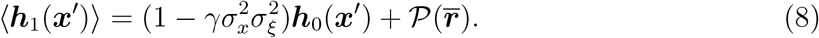

This shows that in our simple linear network model, isotropic variability of the readout weights ***r*** drives compression in all directions within the subspace orthogonal to 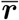. See Fig. S2 for a visualization.

In the above simplified analysis, we assumed a form of variability for ***r*** and directly “imposed” it. In the following, we explore the impact of true variability in ***r*** as driven by SGD (using a batch size of 1 for simplicity). The gradient update for ***r*** in this case is *δ****r*** = −*γe*(***x***)***h***(***x***). This variability affects the hidden representation only after at least two gradient steps. Assuming that the readouts are at equilibrium 〈*δ****r***〉 = **0**, the equation for the updated hidden representation becomes

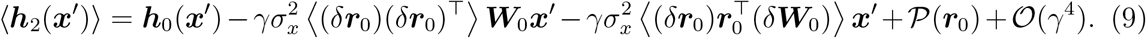

Here *δ* is the update at the first step of SGD, as before. The second term resembles Eq. (6), with 〈(*δ****r***_0_)(*δ****r***_0_)^⊤^〉 replacing 〈***ξξ***^⊤^〉. The third term is an additional cross term that appears due to dependence between the updates to the initial output weights ***r***_0_ and the initial input weights ***W***_0_. For this equation to indicate compression, the norm of the right hand side must be less than the norm of ***h***_0_. While the exact conditions for this to occur are not immediate from the equation, we here outline a set of assumptions that allow it to be analyzed easily, and show that compression occurs in this case. Namely, we assume that ***W***_0_ is a diagonal matrix, ***r***_0_ is proportional to a standard basis vector ***r***_0_ ∝ ***e***_*k*_ for some *k* ∈ {1, 2,…, *N*}, and ***b*** ∝ ***r***_0_. With these assumptions, Eq. (9) decouples into scalar equations

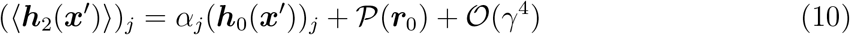

for *j* ≠ *k*, where 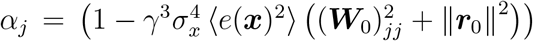 is smaller than 1 for *γ* small enough. Here the 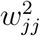 and ‖***r***_0_‖^2^ factors come from the second and third terms of Eq. (9), respectively. This reveals that, for *γ* > 0 small enough and under the simplifying assumptions above, ***h***_2_ is on average reduced in all directions orthogonal to the readout by the noise generated by SGD weight updates.

This equation yields some insights. For one, the effect of compression is of order *γ*^3^, which indicates that the effect may be relatively weak for single-layer networks. For another, the compression depends directly on *e*(***x***)^2^, so that a network that reduces the error very quickly may not compress as much. The term 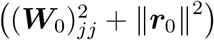 shows dependence of compression on the magnitude of the initial parameters.

This simplified analysis gives intuition for how SGD can lead to representations in which each class of input is mapped into a tightly clustered set. This, in turn, gives rise to an overall dimension for the representation that is close to the number of classes – a dimension that is typically dramatically compressed when compared with the input data itself (see Fig. S3 for simulations showing dependence of dimensionality on the number of classes). To further support our hypothesis of SGD as a mechanism driving compression, in Fig. S5 we compare the dimensionality compression of our RNN model across two cases: using full batches and using a batch size of one. We find that for mean squared error loss the RNN model only compresses dimensionality with minibatching the minibatching. However, both batch sizes cause dimensionality compression in the case of categorical cross entropy loss combined with tanh nonlinearity, indicating that additional mechanisms could be at play.

## CONCLUSIONS AND DISCUSSION

RNNs learn to perform rich operations on their inputs, including expanding and compressing their dimensionality in order solve classification tasks with efficient, compact representations. This behavior relies on the interplay of learning rules with network dynamics. We explore this in two regimes: edge-of-chaos and strongly chaotic dynamics.

Networks in both regimes strongly compress high-dimensional inputs – i.e., those that are initially linearly separable – into distinct sets, one for each class. This behavior also occurs when classifying MNIST (Fig. S4), and has recently been observed in deep networks [12, 49]. We develop mathematical reasoning for why this compression occurs: a linear approximation reveals how SGD itself naturally promotes compression of points belonging to the same class. The compressed representations can arise either from the formation of stable fixed points (for networks initially at the edge of chaos) or relatively low-dimensional chaotic attractors (for networks in the strongly chaotic regime). This has implications for coding: networks that are stabilized by training learn to solve the task for all time while networks that remain chaotic partially “reset” after the evaluation period. The former may lend itself to long-term memory; the latter could be useful in flexibly learning new tasks.

In the case of low-dimensional inputs, we observe differences depending on degree of chaos. Networks initialized on the “edge of chaos” lack an effective mechanism for forming high-dimensional representations, and do not achieve linear separability as a result [13]. Highly chaotic networks, on the other hand, naturally expand dimensionality. The beneficial attributes of dimensionality expansion has been explored in feed-forward models such as kernel learning machines [52] and models of olfactory, cerebellar, gustatory, and visual circuits [3, 5, 8, 39, 43, 46, 57], as well as reservoir computing models where recurrent weights are random and untrained [35].

On the other hand, studies have also pointed out the need to constrain dimensionality to enable generalization [18, 35, 57]. We find that SGD balances the expansion induced by chaos with compression through training, resulting in representations that are both linearly separable as well as compact at the readout time, coinciding with good generalization properties. In the most striking cases (Fig. 2e), strongly chaotic networks initially expand, and then compress dimensionality. In the extreme case of spatially localized inputs constrained to two neurons, the strongly chaotic network typically only expands dimensionality, without subsequently compressing it, seemingly with some cost to generalization; still, the ED remains small relative to the size of the network.

In all of these cases, strongly chaotic networks produce ongoing neural variability, a property shared by biological models of neural circuits [24, 30–32, 38, 44, 45, 51, 59] as well as experimental recordings [40, 45, 56]. Our findings suggest that internally generated variability plays a functional role in neural circuits and can add both biological realism as well as computational power to artificial network models.

A clear need for the future is the consideration of a wider range of tasks (c.f. [14]). Also needed is a more complete study a range of input dimensions as well as higher-dimensional outputs specified by more than two class labels. For a start, we have found that in the case of high-dimensional inputs, the number of class labels strongly modulates the readout dimensionality of the trained RNN, while the number of input clusters does not (Fig. S3). In addition, it remains to extend theoretical arguments for compression induced by SGD to the full nonlinear, recurrent network trained over many samples from multiple classes, and to prove explicitly that this compression results in a reduction of the dimensionality of the network response to inputs. Lastly our results should be generalized to different learning frameworks and models with learning rules other than SGD (see [25] for the case of deep networks trained by an unsupervised learning rule).

Taken together, we find that RNNs learn to balance compression with the natural expansion induced by chaos in a way suitable to the task at hand. These findings invite the further exploration of learning strategies through the lens of modulating dimensionality. The observation that SGD naturally promotes dimensionality compression sheds light on its surprising success in generalizing well (see also [2, 29, 36, 53, 54]). Our observations of compressed *geometry* of learned representations are closely related to prior work on their compressed *information* and allied generalization bounds [53, 54]. Our findings also lend support for the hypothesis that low-dimensional representations in the brain arise naturally from synaptic modifications driven by learning, and highlight a scenario where chaotic variability and salient low-dimensional structure synergistically coexist (cf. [24, 30, 31, 44]).

## METHODS

(See Supplemental Information for more details.)

### Model

We consider a recurrent neural network (RNN) model with one hidden layer trained to perform a delayed classification task (Fig. 1). The equation for the hidden unit activations ***h***_*t*_ is

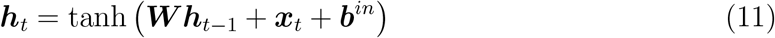

where ***W*** is the matrix of recurrent connection weights, ***x***_*t*_ is the input, and ***b***^*in*^ is a bias term. The network is initialized to have zero activation, ***h***_−1_ = **0**. The output of the network is

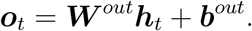

The output ***o*** = ***o***_*t*_eval__ at the evaluation timestep *t*_eval_ = 10 is converted to a scalar error signal via a categorical cross-entropy loss function. This loss is used to update the recurrent weights ***W*** and the output weights ***W***^*out*^ via SGD.

### Inducing Chaos

We initialize ***W*** as ***W*** = (1 − *ε*)***I*** + *ε****J*** where *ε* = .01 ensures smooth dynamics before training. The matrix ***J*** has normally distributed entries that scale in magnitude with a coupling strength parameter *γ*, 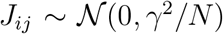. We investigate two dynamical regimes, “edge of chaos” and “strongly chaotic”, with gain strength *γ* = 20 and *γ* = 250, respectively.

## ACKNOWLEDGEMENTS

MF is funded by the National Science Foundation Graduate Research Fellowship under Grant No. DGE-1256082. GL is funded by an NSERC Discovery Grant (RGPIN-2018-04821), an FRQNT Young Investigator Startup Program (2019-NC-253251), and an FRQS Research Scholar Award, Junior 1 (LAJGU0401-253188). ESB acknowledges the support of NSF DMS Grant 1514743. We thank Merav Stern, Dmitri Chklovskii, Ali Weber, Nick Steinmetz and Luca Mazzucato for their insights and suggestions. MF would also like to thank Haim Sompolinsky and SueYeon Chung for their mentorship and inspiration.

## Appendix: Theoretical arguments for compression

Here we show the full steps of the derivation of the compression caused by stochastic gradient descent. The model again is that of a single-layer linear network, with internal neural activations responding to a single input sample ***x*** given by ***h***(***x***) = ***Wx*** + ***b***, and a scalar output by 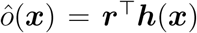. In the following, we assume that ***x*** has zero mean and covariance 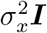. We consider a squared error loss function 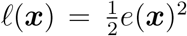, where 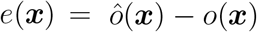 and *o* maps the input ***x*** to its corresponding class label. The gradient descent learning updates over a batch of input samples *B* are:

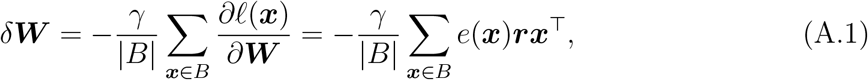

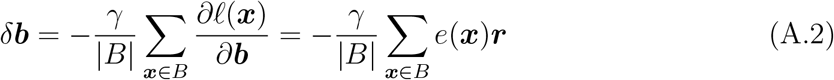

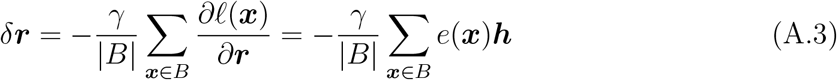

Here *γ* is the *learning rate* for the gradient descent routine and ***rx***^⊤^ is the outer product of ***r*** and ***x***.

Let ***h***_*k*_, and *e_k_* denote the hidden representation and error after *k* steps of gradient descent, respectively, and similarly for parameters ***W***_*k*_, ***b***_*k*_, and ***r***_*k*_. The goal is to compute the hidden representation ***h***_2_(***x***) after two steps of SGD and show that it is compressed when compared with the original hidden representation ***h***_0_(***x***). For SGD with batch size one, on each step of the gradient descent routine an input sample is chosen randomly to use for backpropagating error and updating parameters. Let ***x***_0_ and ***x***_1_ denote the input chosen on the first and second steps of SGD, respectively. As is the usual assumption, we choose ***x***_0_ and ***x***_1_ independently. Let 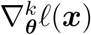 denote the gradient of *ℓ* with respect to parameter ***θ*** evaluated at input ***x*** and at the parameter values at step *k* of SGD (that is, parameters ***W***_*k*_, ***b***_*k*_, and ***r***_*k*_). The parameter updates at the first step are then given by 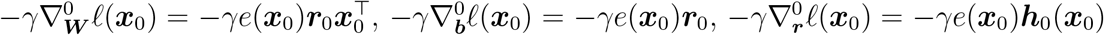 and at the second step by 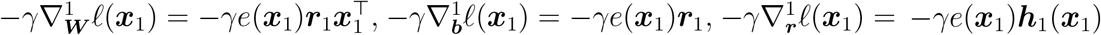. Note that the gradient on the second step depends on the value of the parameters after the first step of gradient descent.

The hidden representation on the second step is ***h***_2_(***x***) = ***W***_2_***x*** + ***b***_2_. The parameters ***W***_2_ and ***b***_2_ are related to their initial values via

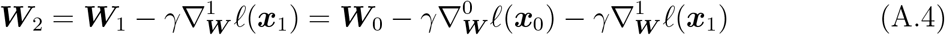

and

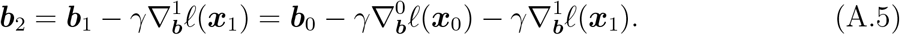

The relationship between ***h***_2_(***x***) and ***h***_0_(***x***) is then given by

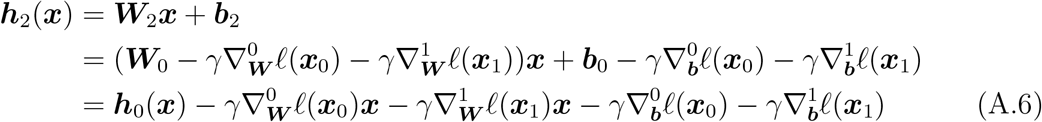

where in the last equation we grouped ***h***_0_(***x***) = ***W***_0_***x*** + ***b***_0_. Our goal is to relate everything in Eq. (A.6) to initial parameter values. This is straightforward for the gradients 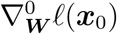 and 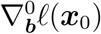: in particular, note that these terms are aligned with ***r***_0_. We will not be needing to consider terms proportional to ***r***_0_ for this analysis, so we collect these terms together. For convenience, suppose that *P* is the orthogonal projector onto the nullspace of ***r***_0_ and let 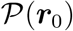 denote any quantity that is mapped to zero by this projector (this notation functions similarly to “big O” notation). Examples of objects that are 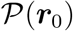 are vectors proportional to ***r***_0_ and matrices of the form ***r***_0_***v***^⊤^. Then ***h***_2_(***x***) can be written

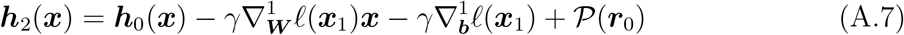

Our next step is to average ***h***_2_(***x***) over choice of ***x***_0_ and ***x***_1_. This results in the expression

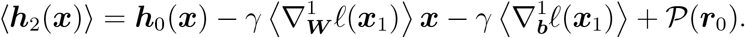

We first assume that the bias is at equilibrium, i.e. 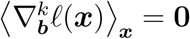. This leaves us with the expression

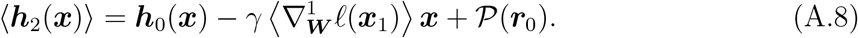

We next focus our attention on computing 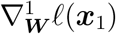. This update is

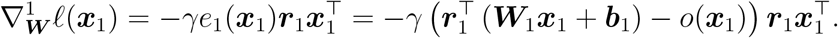

Substituting 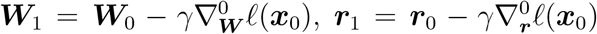 and 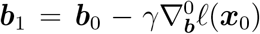 and simplifying results in the expression

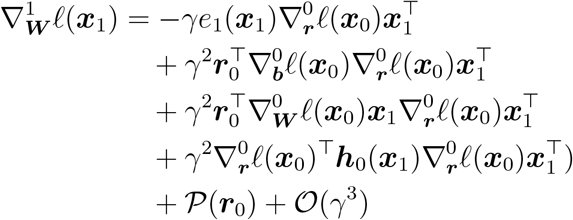

To simplify this expression, we collect terms of order 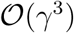 and again collect terms aligned with ***r***_0_. This simplification results in

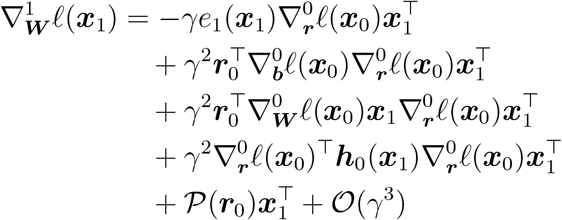

Now we compute the average of this update, 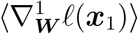. This is

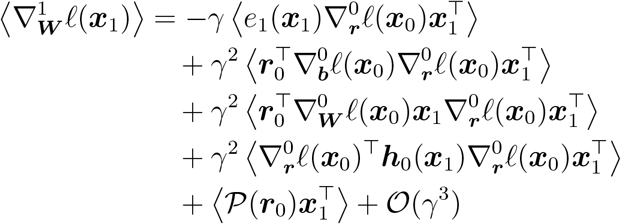

To proceed, we assume that the output weights are at equilibrium, i.e. 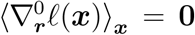. This removes the first term from the expression for 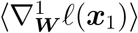. Furthermore, we assume that the input is mean zero and isotropic with variance 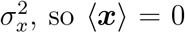, so 〈***x***〉 = 0 and 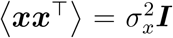. Using these assumptions below, along with some rearranging, we find that the input weight updates satisfy

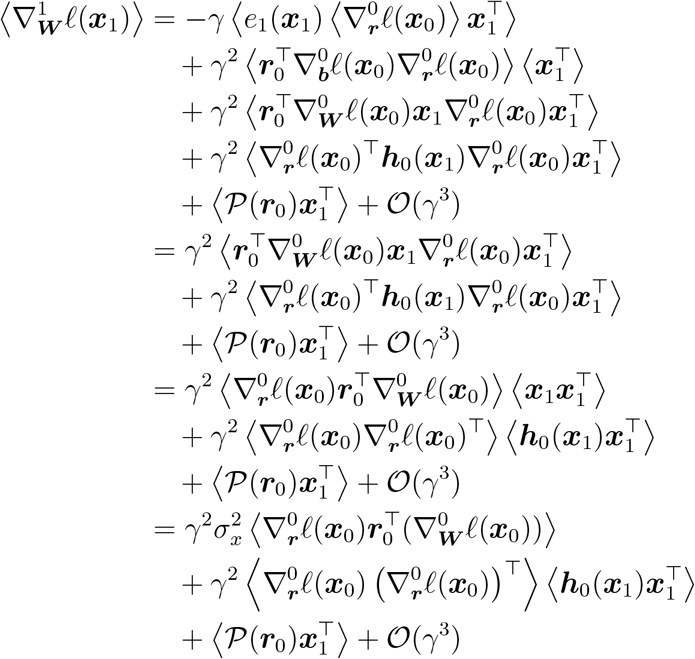

Regarding the term with 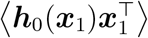, we compute that

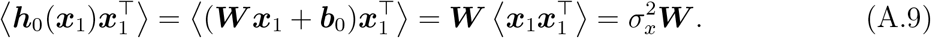

This leaves us with the expression

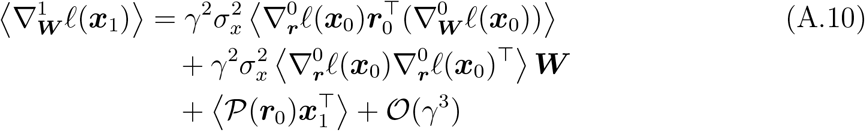

Substituting this expression for 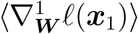 into Eq. (A.8) and simplifying results in

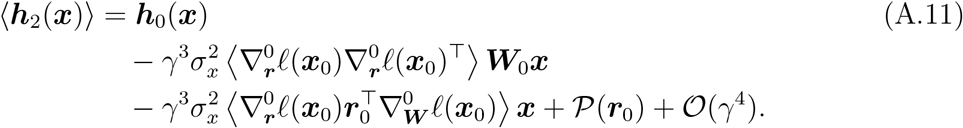

This is a full reduction of the behavior of the representation after two SGD steps in task-irrelevant directions orthogonal to the readouts, up to order 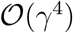, under mild assumptions on the input (mean zero and isotropic) and on the gradients of the output and bias weights (that they are in equilibrium). While a systematic analysis of Eq. (A.11) is desirable, in this work we instead make strong simplifying assumptions that make the equation easy to analyze. The assumptions we make are: (1) ***W***_0_ is diagonal, (2) ***r***_0_ ∝ ***e***_*k*_ is proportional to a standard unit vector ***e***_*k*_, the vector of all zeros except for a one in the *k*th entry, for some *k* ∈ {1, 2,…, *N*}, and (3) ***b***_0_ ∝ ***r***_0_. Let’s assume without loss of generality that ***r***_0_ ∝ ***e***_1_.

We first address the second term in Eq. (A.11) by computing

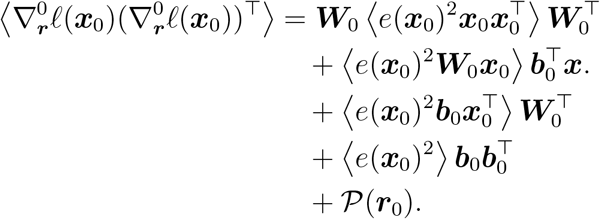

We first deal with the second term of the above expression. Note that by assumptions (1)–(3), *e*(***x***_0_) = ***r***^⊤^(***W***_0_***x***_0_ + ***b***_0_) − *o*(***x***_0_) depends only on the first coordinate of ***x***_0_. Since 〈***x***_0_〉 = **0**, it follows that 〈*e*(***x***_o_)^2^***W***_0_***x***_o_〉 ∝ ***e***_1_ ∝ ***r***_0_. The third and fourth terms are also proportional to ***r***_0_ by assumption (3). Hence

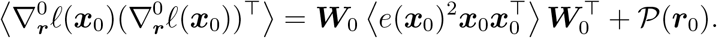

We now finish computing the second term of Eq. (A.11) under assumptions (1)–(3). Using again that *e*(***x***_0_) depends only on the first coordinate of ***x***_0_, as well as the fact that the distinct coordinates of ***x***_0_ are independent, it’s straightforward to show that

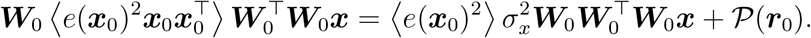

Hence

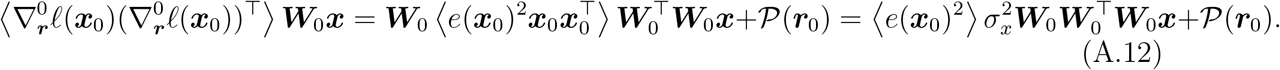

Calculations for the third term of Eq. (A.11) follow a similar flow, resulting in

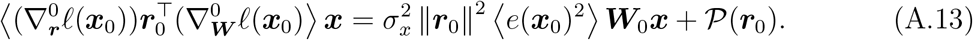

Substituting Eq. (A.13) and Eq. (A.12) into Eq. (A.11) results in

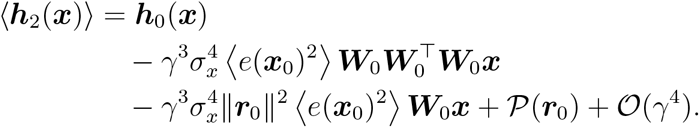

Note that our assumption (3) that ***b***_0_ ∝ ***r***_0_ implies that 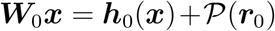. Furthermore, since ***r***_0_ ∝ ***e***_1_ by assumption (2) and ***W***_0_ is diagonal by assumption (1), it follows that 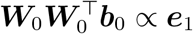 so that 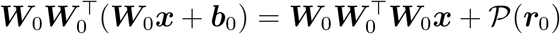. Hence

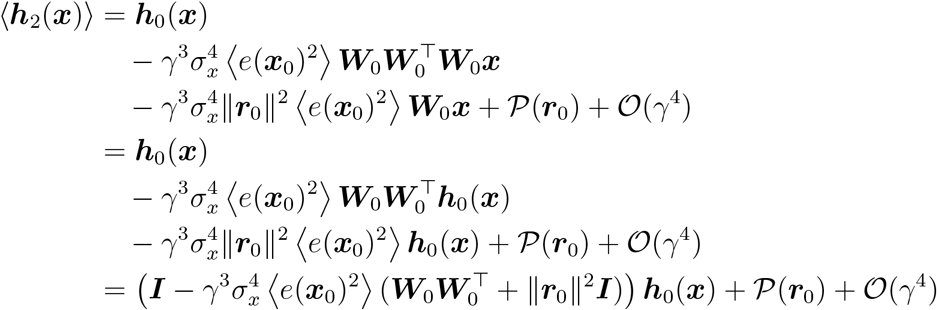

This shows that for *γ* small enough, the hidden representation is scaled by a positive constant less than one in the directions orthogonal to ***r***_0_. This shows that the representation is compressed.

