## Supplemental Information for "Recurrent neural networks learn robust representations by dynamically balancing compression and expansion"

### Supplemental Information (SI)

#### Addition model details

We consider a recurrent neural network (RNN) model with one hidden layer trained to perform a delayed classification task. The equation for the hidden unit activations $\boldsymbol{h}_t$ is

$$\boldsymbol{h}_t = \phi\left(\boldsymbol{W}\boldsymbol{h}_{t-1} + \boldsymbol{x}_t + \boldsymbol{b}^{in}\right) \tag{1}$$

where $\boldsymbol{W}$ is the matrix of recurrent connection weights, $\boldsymbol{x}_t$ is the input, and $\boldsymbol{b}^{in}$ is a bias term. The nonlinearity is $\phi = \tanh$. The network is initialized to have zero activation, $\boldsymbol{h}_{-1} = \boldsymbol{0}$. We write $N$ as the number of hidden units, so for fixed $t$ we have $\boldsymbol{h}_t \in \mathbb{R}^N$.

The output of the network is

$$\boldsymbol{o}_t = \boldsymbol{W}^{out}\boldsymbol{h}_t + \boldsymbol{b}^{out}.$$

The output $\boldsymbol{o} = \boldsymbol{o}_{t_{\text{eval}}}$ at the evaluation timestep $t_{\text{eval}} = 10$ is passed to a categorical cross-entropy loss function to produce a scalar loss signal to be minimized. The equation for this loss is $\text{CCE}(\boldsymbol{o}, k) = \log(p_k)$ where $p_k = \exp(o_k) / \left(\sum_j \exp(o_j)\right)$ represents the probability that $\boldsymbol{o}$ indicates class $k$. This loss is used to update the recurrent weights $\boldsymbol{W}$ and the output weights $\boldsymbol{W}^{out}$ via backpropagation-through-time using the PyTorch 1.0.1 implementation of the RMSProp optimization scheme with a batch size of 10 and learning rate of .001 (loss is summed over the batch). We reduce the learning rate through the training process via a reduce-on-plateau strategy: when the loss fails to decrease by more than $1e-7$ for five epochs in a row, the learning rate is halved. This conservative but fairly standard "reduce only when necessary" approach to the learning rate is meant to help ensure that the quantities we measure over training plateau for reasons other than a vanishing learning rate, while still allowing the network to find the best solution it is able to. The trends and measures analyzed are qualitatively robust to small changes in the learning parameters. Perhaps the parameter with the most significan impact on our results is the initial learning rate. Increasing the learning rate typically increases the amount of the network's compression after training and the degree to which Lyapunov exponents are decreased. In some cases, such as the case of two-dimensional inputs shown to all neurons, doubling the learning rate can have a strong effect in inhibiting the dimensionality expansion of the strongly chaotic network, while halving it is enough to make the compression that follows this expansion considerably fainter. We chose our initial learning rate to roughly maximize the testing accuracy of the networks.

#### Additional task details

The network receives a static input for a number of timesteps (the input period), and is then asked to classify this input after a delay period. In our simulations, the input period is 1 timestep and the delay period is 9 timesteps, after which comes an evaluation period of 1 timestep. Details about the input, delay, and evaluation periods are described in the main text.

The inputs consist of isotropic gaussian clusters distributed randomly through an ambient space of dimension $d$, where each cluster's covariance is the $d \times d$ matrix $\sigma^2\boldsymbol{I}$. The means of the clusters are distributed uniformly at random in a $d$-dimensional hypercube of side length $\ell = 4$ centered at the origin, with a minimum separation $s$ between means being enforced to ensure that clusters do not overlap. Here $\ell$ is chosen so that the average magnitude of the scalar elements of all the input samples is $\sim 1$ (we found this value to roughly maximize the performance of both the edge-of-chaos and strongly chaotic networks). The minimum separation $s$ is chosen so that all points belonging to a cluster fall within a distance $s$ of the mean with a confidence of 99.9999% (i.e. the clusters are non-overlapping with probability close to 1). To form an input sample $\boldsymbol{x}$, we first select a center $\boldsymbol{c} \in \mathcal{C}$ and draw $\boldsymbol{x}$ from the isotropic gaussian distribution centered at

$\boldsymbol{c}$ (which is contained in the ambient space of the input). This is commonly expressed by the notation $\boldsymbol{x} \sim \mathcal{N}(\boldsymbol{c}, \sigma^2 \boldsymbol{I})$. In our case, we choose a standard deviation of $\sigma = 0.02$. To embed these inputs in the $N$-dimensional space of the recurrent units, we multiply the inputs by an orthogonal transformation. In our simulations, we consider the case of 60 clusters, $\#\mathcal{C} = 60$. Each cluster center is randomly associated with one of two class labels (30 clusters for each label), and the training samples drawn from this cluster are assigned this label. Over training, new samples are always drawn, so the network never sees the same training example twice. Test inputs are drawn from the same distribution and have not been seen by the network during training. Our results are not sensitive to small changes in the above parameters.

The input is transformed by a random $N \times d$ orthogonal transformation, where each column has norm $1/\sqrt{d}$ (this transformation preserves the geometry of the inputs). The one exception is input given to two neurons in the network, where the input is transformed by a matrix with 1 in the $(1, 1)$ and $(2, 2)$ positions, and zero everywhere else. The output weights of the network are initialized as a random matrix $\boldsymbol{J'}$ where $J'_{ij} \sim \mathcal{N}(0, .3^2/N)$.

### Inducing chaos

The (intrinsic) dynamics of the network depend primarily on $\boldsymbol{W}$. Strong random coupling between recurrent units leads to chaotic dynamics, whereas weak random coupling results in dynamics that converge onto stable attractors (for large networks these attractors are typically fixed points; see the seminal work by Sompolinsky, Crisanti, and Sommers [7] for an analysis of a rate model network with similar properties). We investigate these dynamical regimes by initializing $\boldsymbol{W}$ as $\boldsymbol{W} = (1 - \varepsilon)\boldsymbol{I} + \varepsilon \boldsymbol{J}$ where $\varepsilon = .01$ sets the timescale of the dynamics to be slow (so that the discrete update Equation (1) produces visually smooth trajectories). The matrix $\boldsymbol{J}$ has normally distributed entries that scale in magnitude with a coupling strength parameter $\gamma$, $J_{ij} \sim \mathcal{N}(0, \gamma^2/N)$. The first regime we consider is at the "edge of chaos" with $\gamma = 20$, where the network is weakly chaotic before training and dynamically stable after training. The second regime we consider is "strongly chaotic" with $\gamma = 250$, where the dynamics are chaotic both before and after training. While our equations are not identical to those analyzed in [7], we find that the trajectories produced and qualitative properties are similar. In particular, the top Lyapunov exponent of our system grows with the gain $\gamma$. In terms of the top Lyapunov exponent, a value of $\gamma = 20$ in our model roughly corresponds to a gain of 2 in the Sompolinsky rate model, while $\gamma = 250$ corresponds to a gain of about 5. The second regime we consider is "strongly chaotic" with $\gamma = 250$, where the dynamics are chaotic both before and after training.

To plot snapshots in time , we compute, independently at each time point, the principal component vectors of the network's responses to the inputs. This means we have a new set of principal component vectors at each time point, and each of these principal vectors is determined only up to sign. To align the points from one time point to the next, we take the signs of the principal vectors that maximize the overlaps between the two time points.

### Measuring chaos

The degree of chaos present in the network is measured through numerical estimates of *Lyapunov exponents*. For a dynamical map $\boldsymbol{h}_{t+1} = f(\boldsymbol{h}_t)$, the Lyapunov exponents $\chi$ are defined as

$$\begin{aligned} \chi(\boldsymbol{h}_0, \delta\boldsymbol{h}_0) &= \lim_{t\to\infty} \frac{1}{t} \ln \frac{\|\delta \boldsymbol{h}_{t+1}\|}{\|\delta \boldsymbol{h}_0\|} \\ &= \lim_{t\to\infty} \frac{1}{t} \ln \|f'(\boldsymbol{h}_t) \cdots f'(\boldsymbol{h}_0)\, \delta \boldsymbol{h}_0\| \,. \end{aligned}$$

where $\boldsymbol{h}_0$ is the initial condition for a trajectory and $\delta\boldsymbol{h}_0$ is a perturbation of $\boldsymbol{h}_0$ in a particular direction. The vector $\delta\boldsymbol{h}_{t+1}$ arises from evolving the perturbation $\delta\boldsymbol{h}_0$ forward in time. The equation measures the rate of expansion/contraction of the neighboring trajectory induced by $\delta\boldsymbol{h}_0$ as the dynamics run forward. For a particular trajectory with initial condition $\boldsymbol{h}_0$, there are $N$ Lyapunov exponents (since the system is $N$-dimensional), depending on choice of $\delta\boldsymbol{h}_0$. The largest exponent $\chi_1$ has particular importance, as $\chi_1 > 0$ indicates chaotic dynamics, and $\chi_1 < 0$ indicates contraction to stable fixed points or stable limit cycles. Since different initial conditions can converge onto different attractors, the Lyapunov exponents can differ with choice of $\boldsymbol{h}_0$. In practice, we find these differences to be small. When we report exponents, we show standard error of the mean over ten randomly chosen initial conditions $\boldsymbol{h}_0$. We use the discrete $\boldsymbol{QR}$ method to numerically compute the exponents [1, 3].

Positive exponents indicate the existence of directions in which nearby dynamical trajectories are pulled apart in time. This persistent pulling apart results in an attractor that has a nonzero "volume" in some subspace: all of the trajectories that collapse to this attractor do not collapse toward the same point, nor a single orbit, and so occupy a nonzero extent of space. This volume is constrained by the negative exponents, which indicate directions in which trajectories are being pulled together. See [4] for details of how this volume is measured, and [5, 6, 2] for investigations of this measure in recurrent networks. If the effective dimensionality of the representation is dominated by a single attractor, we can expect this dimensionality to increase as the number of expansion directions (i.e. the number of positive Lyapunov exponents) increases. This assumption can be violated when there is more than one separate attractor; for instance, different inputs may lead to distinct attractors and thus jointly occupy some subspace of nonzero effective dimension, even if their respective attractors are stable fixed points (which individually have zero volume).

In the case of random (untrained), undriven chaotic networks, one expects that dynamics generally accumulate on a single, connected attractor. The network dynamics return to this attractor after being perturbed by transient inputs. Hence, a strongly chaotic network that is initialized with low-dimensional inputs can be expected to produce dimensionality expansion, as the trajectories return to the higher-dimensional intrinsic chaotic attractor. Note that compression due to negative Lyapunov exponents typically constrains the dimension of the chaotic attractor to be significantly smaller than $N$ [5, 2].

[2] Rainer Engelken and Fred Wolf. Dimensionality and entropy of spontaneous and evoked rate activity. *Bulletin of the American Physical Society*, 2017. Publisher: American Physical Society.

### Additional figures

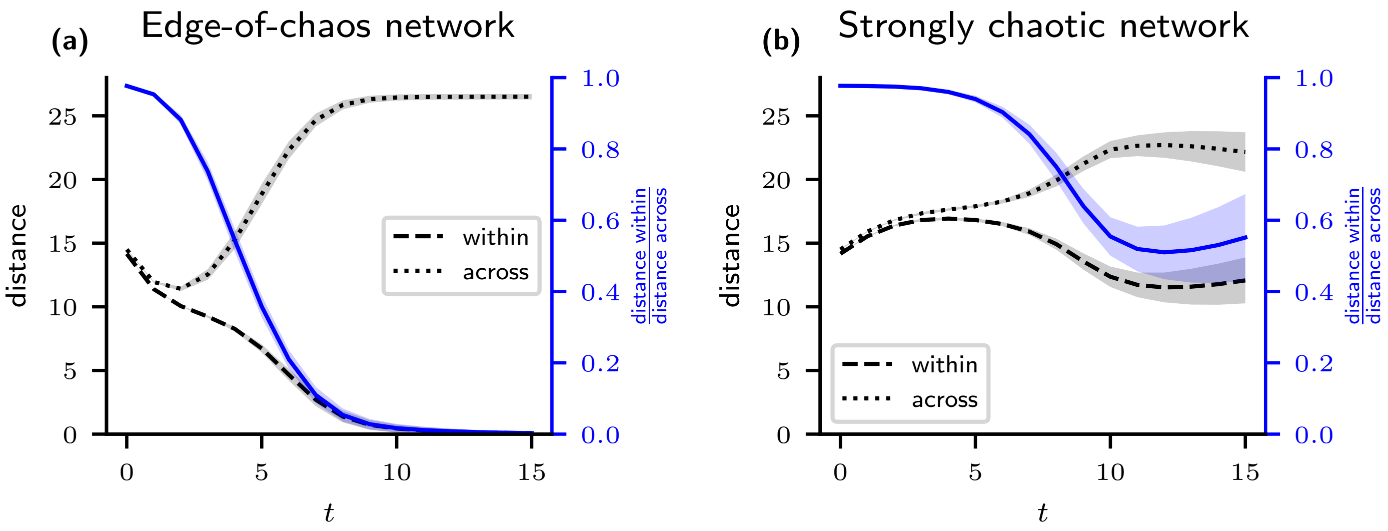


Figure S1: Mean pairwise distance between points belonging to the same class (dashed lines), mean pairwise distance between points belonging to different classes (dotted lines), and the ratio of the first to the second (blue lines and axes). Shaded regions represent 75% probability mass of a normal distribution fit to values of the dependent variables over a sample of 30 network and input realizations, with solid lines indicating medians, for the RNN trained to classify in the high-dimensional inputs case. (a) Edge-of-chaos network as defined in the main text. (b) Strongly chaotic network as defined in the main text.

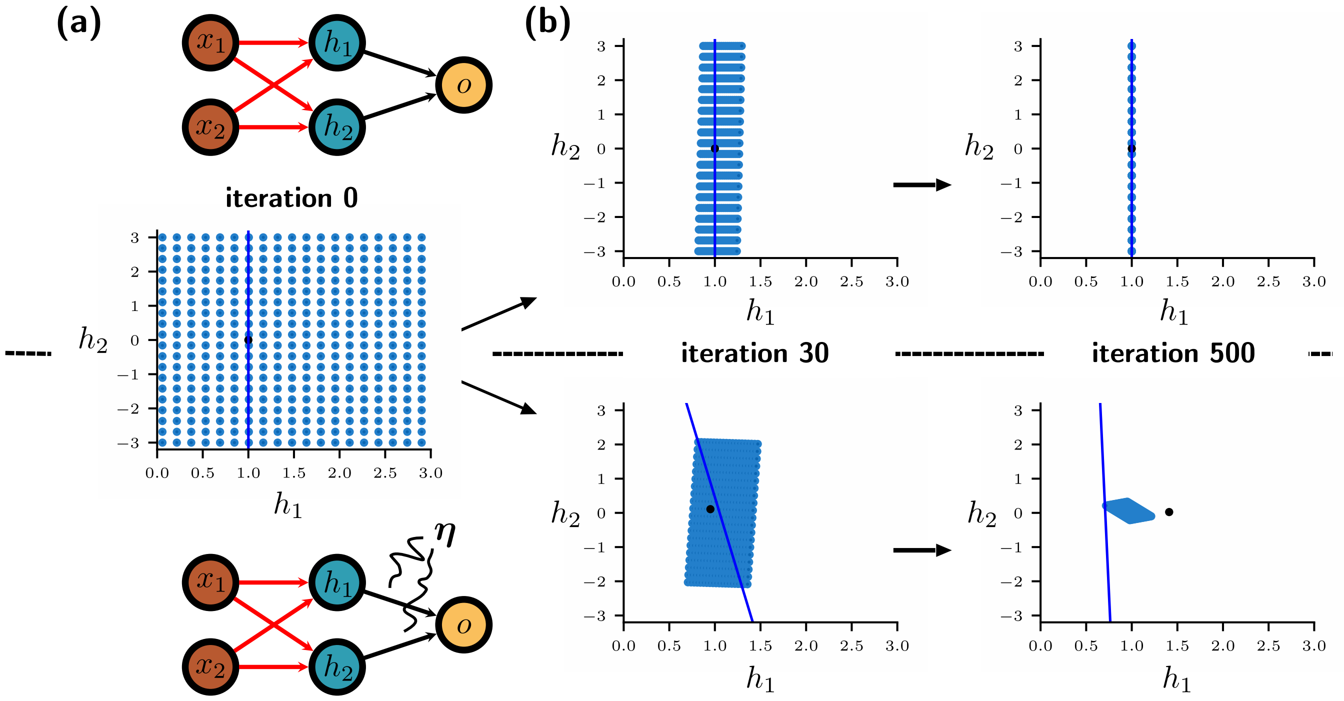


Figure S2: Example of noise in output weights driving compression of the hidden representation in a linear network with two hidden layer units. The equation for the network is $\boldsymbol{h} = \boldsymbol{W}\boldsymbol{x} + \boldsymbol{b}$ with output $o = \boldsymbol{w}^T\boldsymbol{h}$. The input weights (red) are initialized to the $2 \times 2$ identity matrix, and bias is initialized as $(1, 0)$. The inputs are placed on a grid from $x = -1$ to $x = 2$ and from $y = -3$ to $y = 3$ (not shown). Network output $o$ is trained to minimize the squared error loss $0.5(o-1)^2$. Input samples are chosen randomly, and input weights are updated via stochastic gradient descent with batch size 1. (a) Top: Diagram of network where input weights are trained and output weights are fixed. Bottom: Diagram of network where input weights are trained and output weights are normally distributed with mean $(1, 0)$ and covariance $0.05\boldsymbol{I}$. In the figure, $\boldsymbol{\eta}$ represents additive white noise. Middle: hidden unit responses (blue circles) to the inputs before training (iteration 0). Black dot denotes the output weight vector, and the blue line is the affine subspace of points that $w$ maps to 1. (b) Evolution of the hidden layer response to inputs (representation) as input weights are trained. Top: Representation of the network where output weights are fixed. The iteration number denotes the number of training samples that have been used to update the weights. Activations compress to the space orthogonal to $\boldsymbol{w}$, shifted by $(1, 0)$. Bottom: Representation of the network where output weights are randomly drawn at every input sample presentation. Activations compress to a compact, localized space. The direction of compression is both along and orthogonal to $\boldsymbol{w}$.

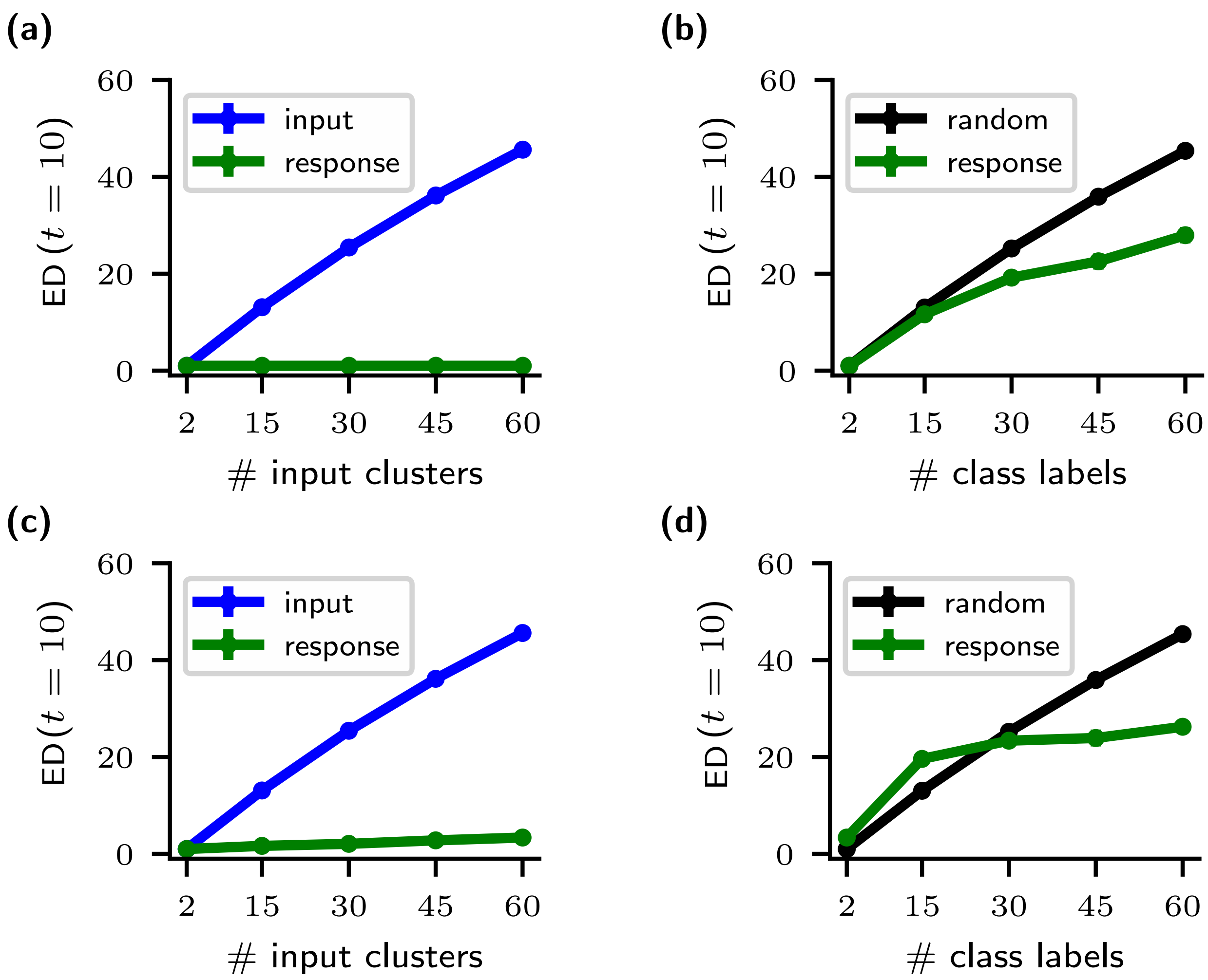


Figure S3: Effective dimensionalities (EDs) of the trained network responses to inputs embedded in an $N$-dimensional ambient space, measured at the evaluation time $t_{eval} = 10$. Error bars denote two standard deviations of three initializations of task and networks (in all panels they are too small to see). (a) Edge-of-chaos networks. Blue: Effective dimensionality of the inputs. Green: effective dimensionality of the network representation as a function of the number of input clusters. Dimensionality remains flat and small. (b) Edge-of-chaos networks. Green: effective dimensionality of the network representation as a function of the number of class labels. Black: Effective dimensionality of points distributed uniformly at random in an $N$-dimensional ball. The number of points drawn is determined by the number of class labels. This is to roughly measure what the effective dimensionality of the network would be if it formed a fixed point for every class label, and distributed these fixed points randomly in space. (c) Strongly chaotic networks. Legend as in (a). (d) Strongly chaotic networks. Legend as in (b).

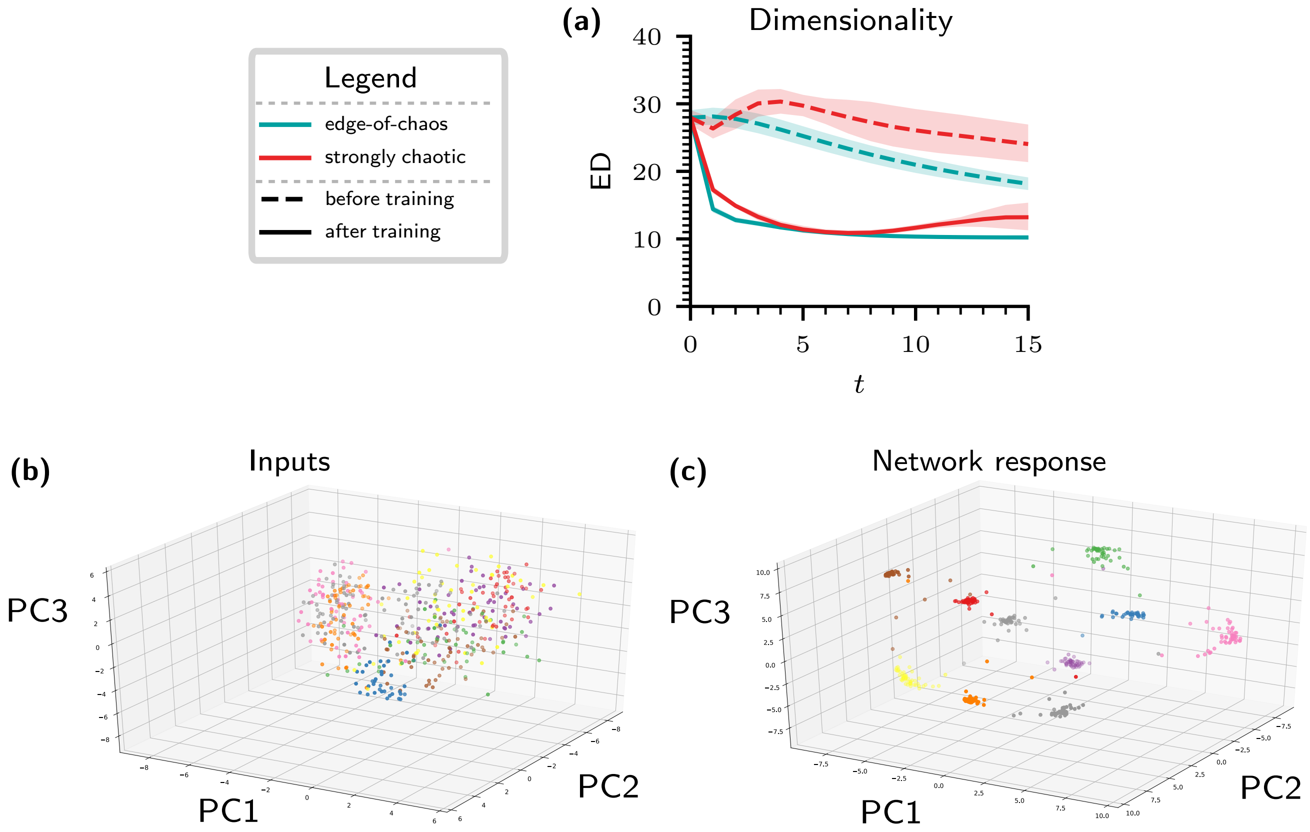


Figure S4: (a) Effective dimensionality (ED) of RNNs trained on the MNIST digit recognition dataset. The ED of the network's responses to test inputs is plotted. Shaded regions indicate 75% probability mass of gamma distributions fit to values of the dependent variables over a sample of 10 network and input realizations, with solid lines indicating medians. After training, dimensionality compresses down to a value by $t = 10$ that roughly matches the number of class labels (10). This compression is similar to that seen in Fig. 2 of the main text. (b) Projection onto the top three principal components of MNIST test data. Colors indicate true class label (i.e. digit identity). (c) Projection onto the top three principal components of the edge-of-chaos recurrent network's responses to the inputs in (b) after training, at the evaluation time $t = 10$. Colors indicate true class label as in (b). The network forms a localized cluster for each digit.

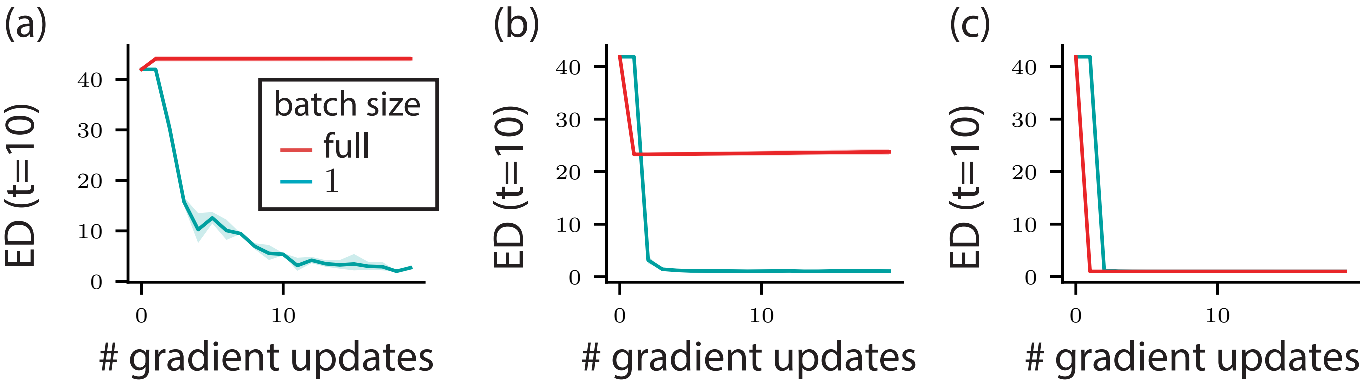


Figure S5: Effective Dimensionality (ED) of the hidden representation at $t = 10$ evaluated through training, comparing the effects of minibatching (blue-green) and full batches (red). Note that the # of gradient updates implies different numbers of training samples for different batch sizes: for a batch size of one, each gradient update follows feeding one input sample to the network, while for a full batch size each gradient update follows only after feeding 200 input samples to the network. In both cases the learning rate is the same. (a) Result for a linear recurrent network trained via mean squared error loss. Only the minibatch updates result in dimensionality compression. (b) Result for a recurrent network with a tanh nonlinearity, trained using a mean squared error loss. The full batch network compresses dimensionality, but much less than the minibatch network. (c) Result for a recurrent network with a tanh nonlinearity, trained using a categorical cross entropy loss. In this case both networks compress dimensionality.
